## Supplementary Materials for "Electronic “photoreceptors” enable prosthetic vision with acuity matching the natural resolution in rats"

### 1 Circuit Dynamics of the Implant

Monopolar electrode arrays with photovoltaic pixels arranged in hexagonal grids are used in this study. Each pixel consists of a photodiode and an active electrode, and the return electrodes of all the pixels are connected to a large SIROF ring in the periphery of the implant.[1] Since electric potentials generated by the electrodes in electrolyte are affecting each other, the circuit dynamics of the full array must be calculated taking into account this coupling.

#### *1.1 Cross-Pixel Impedance and the Access Resistance Matrix*

Consider the electric field where pixel $m$ injects a current of $I_m$ into the electrolyte while all other pixels are idle, as illustrated in Fig S1A. The electric current $I_m$ elevates the potential at the active electrode of pixel $m$ on the electrolyte side by $\Delta V_{m,m}$, as well as the potentials at the neighboring pixel $m-1$ and pixel $m+1$ by $\Delta V_{m-1,m}$ and $\Delta V_{m+1,m}$, respectively. More generally, we denote the potential elevation at the active electrode of pixel $l$ resulting from the current injection at pixel $m$ by $\Delta V_{l,m}$, and define the cross-pixel resistance as

$$R_{l,m} := \frac{\Delta V_{l,m}}{I_m}, \tag{1}$$

where $I_m$ is the current injected by pixel $m$ individually, and the convention is chosen such that $I_m > 0$ for anodal current. Note that $R_{m,m}$ is the conventional definition of the access resistance of the pixel $m$.

Since the volume conduction in electrolyte is considered linear in the range of our study, based on the principle of superposition, when pixels inject current simultaneously, the electric potential

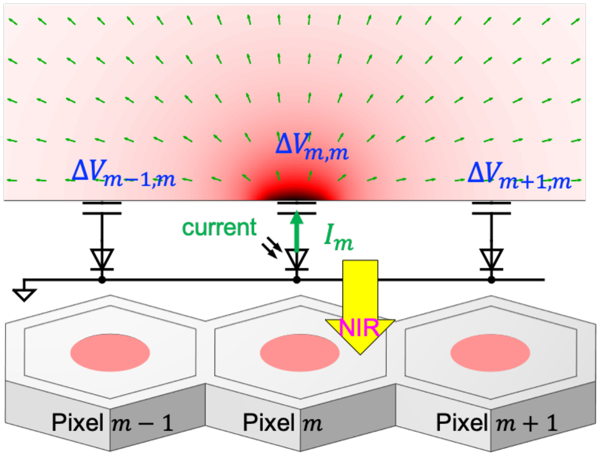


**Fig S1** Schematic illustration of the spatial coupling of the electric potential in electrolyte. The center pixel $m$ is illuminated and injects current $I_m$, while the two neighboring pixels are kept in the dark, without the current injection.

$V_l$ at pixel $l$ is the sum of contributions from all pixels, weighted by their respective currents:

$$V_l = \sum_m R_{l,m} I_m. \tag{2}$$

Let $N$ be the number of pixels on the implant (N=821 in our case of $40\,\mu\mathrm{m}$ pixels). We define the potential vector $\boldsymbol{V} := [V_1, V_2, \cdots, V_N]^\mathsf{T}$, the current vector $\boldsymbol{I} := [I_1, I_2, \cdots, I_N]^\mathsf{T}$, and the access resistance matrix

$$\mathbf{R}_\mathrm{a} := \begin{bmatrix} R_{1,1} & R_{1,2} & \cdots & R_{1,N} \\ R_{2,1} & R_{2,2} & & \\ \vdots & & \ddots & R_{N-1,N} \\ R_{N,1} & & R_{N,N-1} & R_{N,N} \end{bmatrix}. \tag{3}$$

By Equation (2), we have

$$\boldsymbol{V} = \mathbf{R}_\mathrm{a}\boldsymbol{I}. \tag{4}$$

$\mathbf{R}_\mathrm{a}$ is numerically computed by the finite-element method with a parametric sweep in COMSOL Multiphysics 5.6.

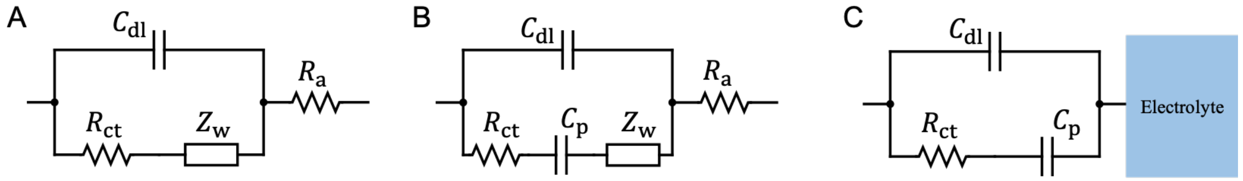


**Fig S2** **A**: The classic Randles circuit model. **B** Circuit model of a pseudocapacitive electrode-electrolyte interface, with a pseudocapacitor in the lower arm of **A**. **C**: Simplified pseudocapacitive model when the interface is not mass-transfer limited and the ohmic potential drop in electrolyte is separately characterized by the access resistance matrix shown in Figure S3.

#### *1.2 The Electrode-Electrolyte Interface*

The pseudocapacitive electrode-electrolyte interface of the SIROF electrode can be modeled with the Randles circuit (Fig S2A) comprising the access resistance $R_a$, the double layer capacitance $C_{dl}$, and the Faradaic arm[2]including the charge transfer resistance $R_{ct}$, the Warburg impedance $Z_w$ and a pseudocapacitor $C_p$ , as shown in Figure S2B. The access resistance has been characterized in its multi-dimensional form in Section 1.1. When the electrochemical processes are not mass-transfer limited, such as SIROF electrode with a typical current density for neural stimulation,[3] the effect of the Warburg impedance is negligible, and hence the interface model can be simplified to Fig S2C.

For the experiments *in vitro* with 1:10 diluted PBS solution (to match the retinal resistivity), the circuit model is fit to the waveforms recorded by the micro-pipette, and yields $R_{ct} = 0.75\,\Omega\,\mathrm{cm}^2$, $C_p = 8\,\mathrm{mF/cm}^2$ and $C_{dl} = 0.3\,\mathrm{mF/cm}^2$. The *in-vivo* circuit model is obtained by fitting to the electric potential measured on the rat cornea when projecting NIR pulses onto the implant in the subretinal space at various frequencies, yielding $R_{ct} = 5\,\Omega\,\mathrm{cm}^2$, $C_p = 9\,\mathrm{mF/cm}^2$ and $C_{dl} = 1\,\mathrm{mF/cm}^2$. The volume conduction model of the rat eye and the head is described in Section 2. The *in-vitro* double-layer capacitance is smaller than the *in-vivo* value approximately by a factor of 3, because 10-fold diluted PBS was used in the *in-vitro* experiment, and the Debye length scales with the square root of the ion concentration. The difference between the *in-vivo* and *in-vitro* pseudocapacitive parameters is likely due to the difference in chemical compositions of the electrolytes.

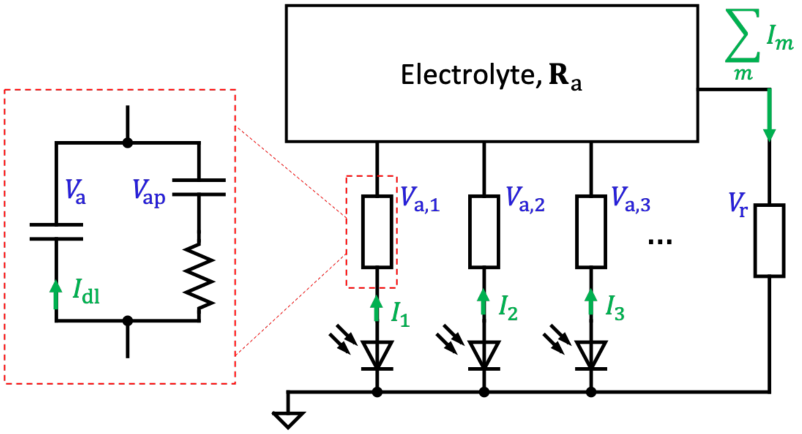


**Fig S3** Circuit diagram of the array of photovoltaic pixels in electrolyte. The small blocks represent the lumped circuit model of the electrode-electrolyte interface shown in Fig S2B, not to be confused with the IEC-style resistors.

#### *1.3 Dynamics of the Photovoltaic Array*

Each pixel on the implant is equipped with a photodiode converting NIR light into anodal current injected into electrolyte through the active electrode. Because of the spatial coupling of the electric potentials described in Section 1.1, the electrolyte is modeled as a multi-port component, to which all electrodes are connected via the interface circuit model described in Section 1.2 and illustrated in Figure S3.

Photocurrent of the pixel $m$ relates to the local irradiance $S_m$ by $I_{\text{pho},m} = \eta S_m$, where $\eta$ is the light-to-current conversion factor.[1] The current injection by pixel $m$ equals the difference between the photocurrent and the forward current of the pn junction:

$$I_m = I_{\text{pho},m} - I_0 \left( \exp\left( \frac{V_{\text{diode},m} e}{n k_\text{B} T} \right) - 1 \right), \tag{5}$$

where $I_0$ is the saturation current, $V_{\text{diode},m}$ the forward-bias voltage of the pn junction, $e$ the elementary charge, $n$ the ideality factor of the diode, $k_\text{B}$ the Boltzmann constant, and $T$ the temperature.[4] For any given irradiance, let $I_m$ be a function $D$ of $V_{\text{diode},m}$ defined by Equation (5). Although the dark *I*-*V* characteristics of our diodes can be adequately approximated by $I_0 = 0.3\,\text{pA}$ and $n = 1.5$,[1] we compute $D(V_{\text{diode},m})$ by interpolating the measured *I*-*V* curve for the best accuracy.

Let $V_{\text{a},m}$ be the potential drop across the electrode-electrolyte interface at the active electrode of the pixel $m$, and $V_\text{r}$ the interface potential at the return electrode. By the Kirchhoff's Voltage Law (KVL), we have, for all $m \in \{1, 2, \cdots N\}$,

$$V_{\text{a},m} + \sum_l R_{m,l} I_l - V_\text{r} - V_{\text{diode},m} = 0. \tag{6}$$

We define $\boldsymbol{V}_\text{a} := [V_{\text{a},1}, V_{\text{a},2}, \cdots, V_{\text{a},N}]^\intercal$, $\boldsymbol{V}_\text{diode} := [V_{\text{diode},1}, V_{\text{diode},2}, \cdots, V_{\text{diode},N}]^\intercal$ and extend the definition of $D$ to operate element-wise on $\boldsymbol{I}$, such that $\boldsymbol{I} = D(\boldsymbol{V}_\text{diode})$. The multi-dimensional form of Equation (6) is

$$\boldsymbol{V}_\text{a} + \mathbf{R}_\text{a} D(\boldsymbol{V}_\text{diode}) - V_\text{r} - \boldsymbol{V}_\text{diode} = 0. \tag{7}$$

To compute the electrode dynamics of the electrode-electrolyte interface, we denote $V_{\text{ap},m}$ the pseudocapacitive voltage at the active electrode of pixel $m$, and $I_{\text{dl},m}$ the charging rate of the double layer, as shown in the dashed red box in Fig S3. For all $m \in \{1, 2, \cdots N\}$, we have

$$C_\text{dl} \frac{d}{dt} V_{\text{dl},m} = I_{\text{dl},m} \tag{8}$$

for the dynamics of the double-layer capacitance,

$$C_\text{p} \frac{d}{dt} V_{\text{ap},m} = I_m - I_{\text{dl},m} \tag{9}$$

for the dynamics of the pseudocapacitance, and, by KVL,

$$R_\text{ct}(I_m - I_{\text{dl},m}) + V_{\text{ap},m} = V_{\text{a},m}. \tag{10}$$

Similarly, we can derive the dynamics for the return electrode, where the current is a sum of the currents of all active electrodes. Equations (7) to (10) describe the interrelated dynamics of all the pixels.

### 2 Model for the Volume Conduction in the Rat Eye and Head

A 3-dimensional model of the rat eye was constructed in Solidworks 2020 (Dassault Systemes Solidworks Corp., MA), similarly to the model by Selner *et al.* for electroretinogram,[5] except for the conductivity and the thickness of the retina, which were modified to $1\,\mathrm{mS\,cm^{-1}}$ and $100\,\mathrm{\mu m}$,[6] respectively, to match the degenerate retina.

The geometry of an anatomically realistic rat head was obtained from `cgtrader.com` (CG-Trader, NY), scaled and combined with the eye for volume-conduction modeling. Conductivity of the brain tissue, $5\,\mathrm{mS\,cm^{-1}}$,[7] was assigned to the head as an approximation of the average conductance. To compute the *in-vivo* elementary electric fields, the geometry was imported to COMSOL Multiphysics 5.6 (COMSOL, Inc., Sweden) for modeling the volume conduction of electric current, with the conductivity of various anatomical structures listed in Table S1. The structures are depicted in Figure S4B, showing a cross-sectional diagram of the ocular and orbital anatomy, with an implant in the subretinal space. Insulation of the eye from the rest of the body by orbital tissues, such as the adipose fat and bone, is necessary to achieve realistic corneal potential in the model, and its amplitude varies with conductivity of these tissues.

Figure S4A shows an elementary field in the rat head generated by a center pixel injecting $1\,\mathrm{\mu A}$ of current. Although the anatomically accurate model can be used to compute the electric fields, its complex shape compromises the reliability of the meshing grid and its incorporation with the model eye. Therefore, the model is prone to computational errors, especially when interacting with other entities, such as ocular tissue layers and the implant, defined in COMSOL, hence complicating further adjustments of the geometry. Since the electric field is confined within several millimeters around the eye, as shown in S4A, the complex anatomically accurate model of the rat head is unnecessary, and we simplify it by replacing the rat head with a cylinder of $8\,\mathrm{cm}$ in height and $4\,\mathrm{cm}$ in radius, while keeping all the ocular and orbital tissues the same. The diagram in Figure S4B shows the anatomy of the simplified model, in which the bone and the skin extend to the cylindrical side surface (not shown).

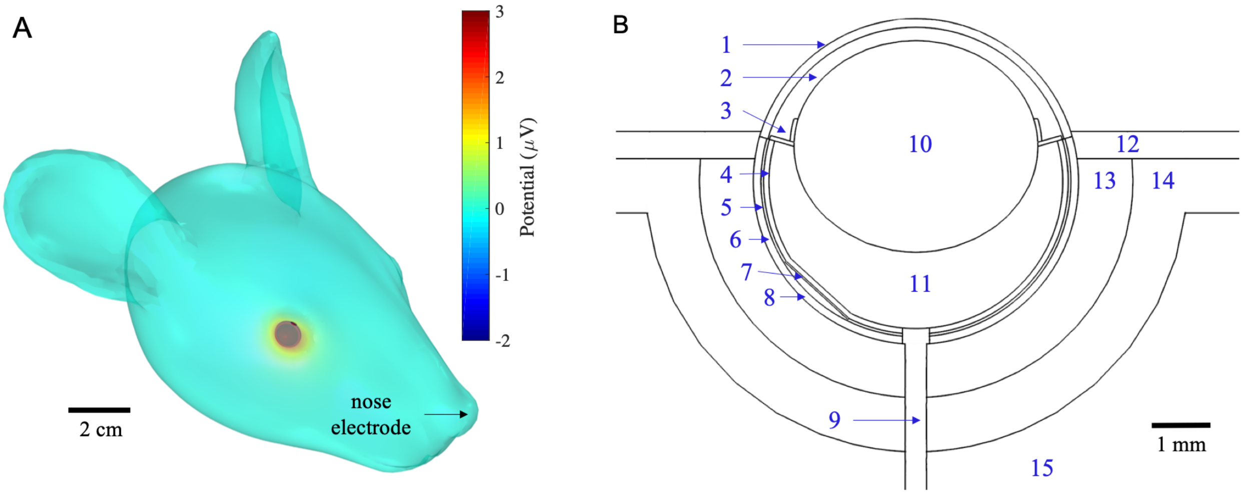


**Fig S4** **A**: Elementary electric field in the anatomically realistic model of the rat head, generated by the subretinal implant with $40\,\mu m$ pixels injecting $1\,\mu A$ of current from an active electrode at its center. **B**: Cross-sectional diagram of the ocular and orbital anatomy in the simplified model with a cylindrical head. Keys to the indices are in Table S1.

| **Index** | **Structure** | **Conductivity** ($mS\,cm^{-1}$) | **Reference** |
|---|---|---|---|
| **1** | Cornea | 4.22 | Selner et al.[5] |
| **2** | Anterior chamber | 15.0 | Selner et al.[5] |
| **3** | Iris | 2.67 | Selner et al.[5] |
| **4** | Retina | 1.00 | Flores et al.[6] |
| **5** | RPE | 0.021 | Selner et al.[5] |
| **6** | Choroid | 2.78 | Selner et al.[5] |
| **7** | Implant | n/a | |
| **8** | Sclera | 5.03 | Selner et al.[5] |
| **9** | Optic nerve | 0.28 | Selner et al.[5] |
| **10** | Lens | 3.22 | Selner et al.[5] |
| **11** | Vitreous | 15.0 | Selner et al.[5] |
| **12** | Skin | 3.04 | Faes et al.[8] |
| **13** | Adipose fat | 0.26 | Faes et al.[8] |
| **14** | Bone | 0.091 | Balmer et al.[9] |
| **15** | Intracranial tissues | 2.00 | Koessler et al.[7] |

**Table S1** Structures labeled in Figure S4B, the conductivity values assigned, and the corresponding references.

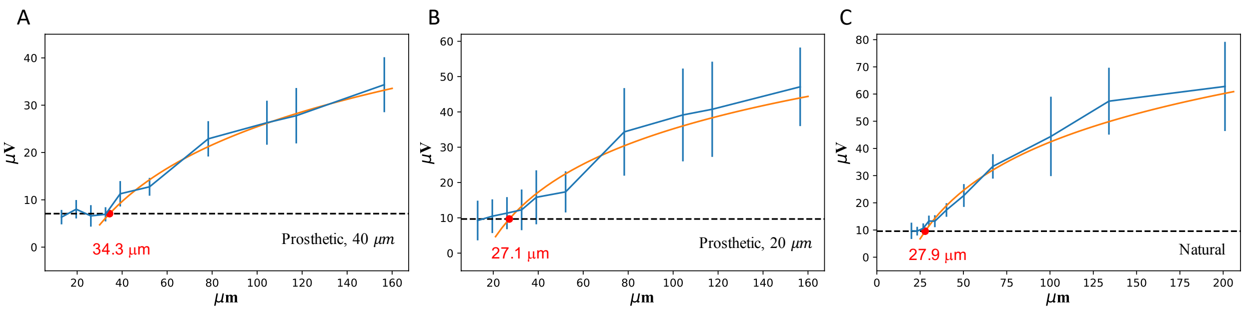


**Fig S5** Prosthetic and natural visual acuity measurements in rats. A-C) VEP amplitude as a function of the grating bar width for prosthetic vision with implants of 40 $\mu$m pixels (A), 20 $\mu$m pixels (B), and for natural vision (C). The red fitting line is the same logarithm function as in Figure 7.

*References*


1 T. W. Huang, T. I. Kamins, Z. C. Chen, *et al.*, "Vertical-junction photodiodes for smaller pixels in retinal prostheses," *Journal of Neural Engineering* **18**(3), 036015 (2021).

2 M. S. Halper and J. C. Ellenbogen, "Supercapacitors: A brief overview," *The MITRE Corporation, McLean, Virginia, USA* **1** (2006).

3 Z. Chen, L. Ryzhik, and D. Palanker, "Current distribution on capacitive electrode-electrolyte interfaces," *Physical Review Applied* **13**(1), 014004 (2020).

4 D. Boinagrov, X. Lei, G. Goetz, *et al.*, "Photovoltaic pixels for neural stimulation: circuit models and performance," *IEEE transactions on biomedical circuits and systems* **10**(1), 85–97 (2015).

5 A. N. Selner, Z. Derafshi, B. E. Kunzer, *et al.*, "Three-dimensional model of electroretinogram field potentials in the rat eye," *IEEE Transactions on Biomedical Engineering* **65**(12), 2781–2789 (2018).

6 T. Flores, T. Huang, M. Bhuckory, *et al.*, "Honeycomb-shaped electro-neural interface enables cellular-scale pixels in subretinal prosthesis," *Scientific reports* **9**(1), 1–12 (2019).

7 L. Koessler, S. Colnat-Coulbois, T. Cecchin, *et al.*, "In-vivo measurements of human brain tissue conductivity using focal electrical current injection through intracerebral multicontact electrodes," *Human brain mapping* **38**(2), 974–986 (2017).

8 T. Faes, H. Van Der Meij, J. De Munck, *et al.*, "The electric resistivity of human tissues (100 hz-10 mhz): a meta-analysis of review studies," *Physiological measurement* **20**(4), R1 (1999).

9 T. W. Balmer, S. Vesztergom, P. Broekmann, *et al.*, "Characterization of the electrical conductivity of bone and its correlation to osseous structure," *Scientific reports* **8**(1), 1–8 (2018).
